# DNA segment of African Swine Fever Virus first detected in hard ticks from sheep and bovine

**DOI:** 10.1101/485060

**Authors:** Ze Chen, Xiaofeng Xu, Xiaojun Yang, Weihao Dou, Xiufeng Jin, Haishuo Ji, Guangyuan Liu, Jianxun Luo, Hong Yin, Shan Gao

## Abstract

In this study, we aimed to detect viruses in hard ticks using the small RNA sequencing based method. A 235-bp DNA segment was detected in *Dermacentor nuttalli* (hard ticks) and *D. silvarum* (hard ticks) from sheep and bovine, respectively. The detected 235-bp segment had an identity of 99% to a 235-bp DNA segment of African Swine Fever Virus (ASFV) and contained three single nucleotide mutations (C38T, C76T and A108C). C38T, resulting in an single amino acid mutation G66D, suggests the existence of a new ASFV strain, which is different from all reported ASFV strains in NCBI GenBank database. These results also suggest that ASFV could have a wide range of hosts or vectors, beyond the well known Suidae family and soft ticks. Our findings pave the way toward further studies of ASFV transmission and development of prevention and control measures.

## Introduction

African Swine Fever Virus (ASFV) is a large (~ 190 Kbp), double-stranded DNA virus with a linear genome containing at least 150 genes. ASFV causes African Swine Fever (ASF), a highly contagious viral disease of swine. This disease results in high mortality, approaching 100% [1]. ASF first broke out in Kenya in 1921 [2] and remained restricted to Africa until 1957, when it was reported in Portugal. In the following years, ASF spread further geographically and caused economic losses on the swine industry. In 2018, ASF was reported in Liaoning province of China and then spread to more than 20 provinces. Although ASFV can be fast detected using PCR with specific primers [3], the understanding of ASFV vectors are still incomplete [4].

ASFV infects only members of the Suidae family including domestic pigs, warthogs and bushpigs [5]. It is well accepted that only soft ticks belonging to the genus *Ornithodoros* are potential biological vectors of ASFV. To the best of our knowledge, ASFV has not been reported to be detected in *Dermacentor nuttalli, D. silvarum* and *Amblyomma testudinarium*, which are three species belonging to Ixodidae (hard ticks). *Dermacentor* ticks share similarities in the range of hosts as *Ornithodoros* ticks, but *Dermacentor* ticks have higher mobility and a wider range of geographic distribution. Compared to *Ornithodoros* and other *Ixodidae* ticks, *A. testudinarium* ticks have a wider range of geographic distribution in south of China, where ASF has been reported in 2018 but *Ornithodoros* occurrences have not been documented. *A. testudinarium* ticks have a wider range of hosts covering all members in the Suidae family. In addition, *Dermacentor* and *Amblyomma* ticks produce ten times eggs than *Ornithodoros* ticks. The viruses transmitted by *D. nuttalli, D. silvarum* and *A. testudinarium* are under-estimated as those tramsmitted by insects, based on our previous studies using high-throughput sequencing [6].

Small RNA sequencing (small RNA-seq or sRNA-seq) is used to obtain thousands of short RNA sequences with lengths that are usually less than 50 bp. sRNA-seq has been successful used for virus detection in plants [7], invertebrates and human [8]. In 2016, an automated bioinformatics pipeline VirusDetect has been reported to facilitate the large-scale virus detection using sRNA-seq [6]. This study aimed to detect viruses in *D. nuttalli*, *D. silvarum* and *A. testudinarium* using the sRNA-seq based method. As a unexpectedly result, VirusDetect reported the existence of ASFV in *D. nuttalli* from sheep and bovine. To confirm this result, a 235-bp DNA segment of ASFV was detected in *D. nuttalli, D. silvarum* and sheep, but not in *A. testudinarium* and bovine. Although we only obtained two 235-bp DNA sequences from *D. silvarum* and sheep, respectively, our study still proved the existence of ASFV in *D. nuttalli, D. silvarum* and sheep,

## Results

After data cleaning and quality control, 13,496,191, 25,194,632, 37,888,277, 12,302,335 and 15,077,054 cleaned reads were used to detect viruses in *A. testudinarium* adults, *D. nuttalli* adults, *D. nuttalli* adults (replicate), *D. nuttalli* larvae and *D. nuttalli* nymphs (**Materials and Methods**). VirusDetect reported the existence of ASFV in all four *D. nuttalli* samples, but not in *A. testudinarium* adults. Aligning the cleaned reads to the ASFV reference genome (GenBank: AY261365.1), the mapping rates of four *D. nuttalli* samples reached 0.06% (15,585/25,194,632), 0.06% (23,330/37,888,277), 0.08% (10,241/12,302,335) and 0.08% (12,807/15,077,054), which were significantly higher than the mapping rate 0.01% (905/13,496,191) of *A. testudinarium* adults. The length distribution of virus derived small RNAs (vsRNAs) in four *D. nuttalli* samples concentrated in 15 - 19 bp rather than 21 - 24 bp. The length distribution of vsRNAs was different from those in plant [9] and other invertebrate vsRNAs [10].

To validate the existence of ASFV, we used PCR with specific primers to amplify a 235-bp ASFV segment using total RNA of the *D. nuttalli* adults (**Materials and Methods**), from which ASFV had been detected using sRNA-seq. The gel electrophoresis result showed a clear 235-bp band as we expected, but Sanger sequencing of it failed due to low DNA concentration. To confirm these results, we used PCR to detect this segment in *D. silvarum* ticks, sheep and bovine collected in Xinjiang province of China. The gel electrophoresis results showed the 235-bp bands appeared in 33 and 12, out of 80 *D. silvarum* samples and 100 sheep blood samples, respectively (**Figure 1A**). Two 235-bp DNA sequences were obtained using Sanger sequencing from *D. silvarum* and sheep, respectively to confirm that these 235-bp bands are the 235-bp ASFV segment (**Figure 1B**). In addition, the gel electrophoresis results showed 560-bp bands in 41 out of 100 bovine samples. Then, we obtained two 560-bp sequences using Sanger sequencing. Using blast tools, the results showed that these 560-bp bands could be segments from the bovine genome.

**Figure 1.**
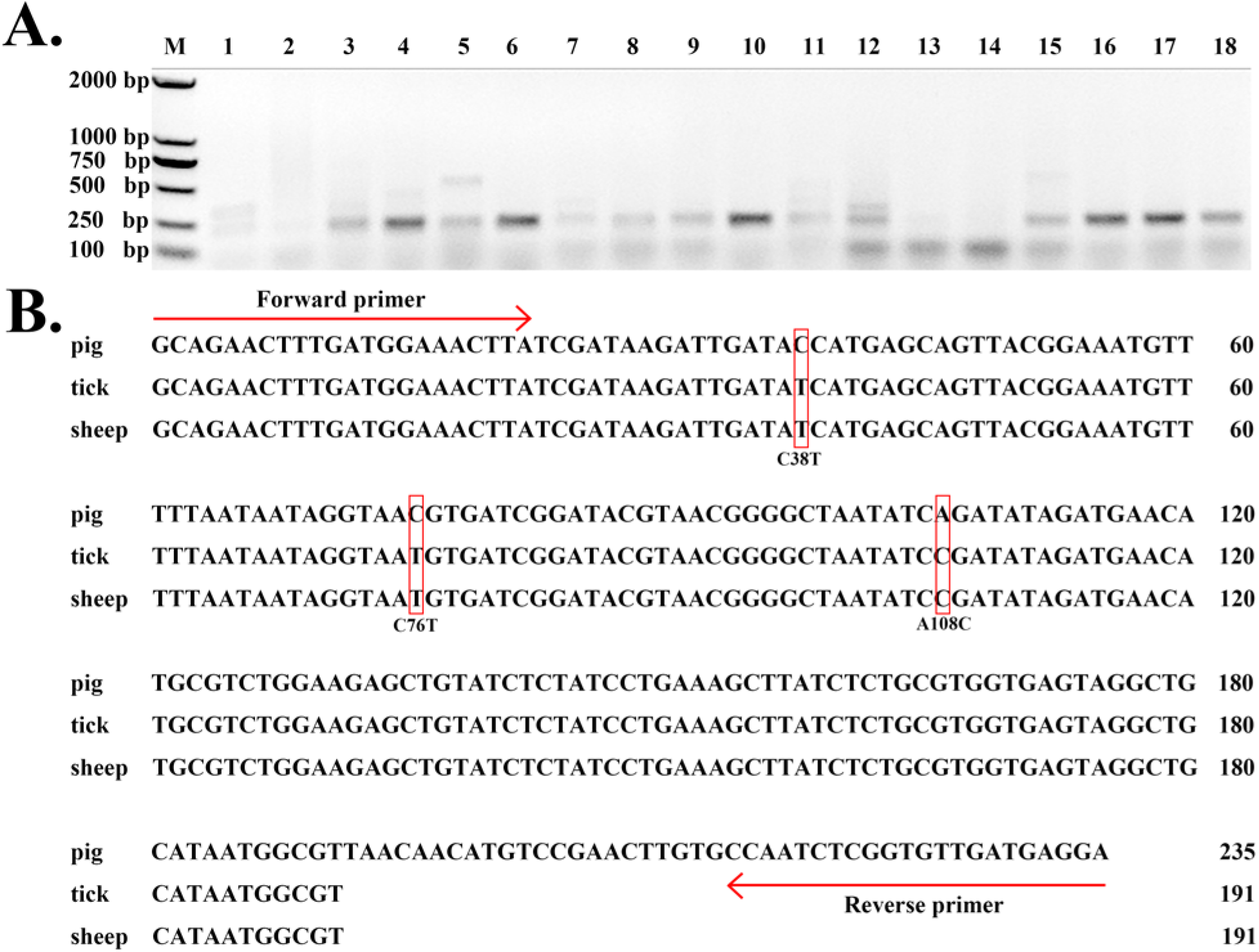
235-bp DNA segments of ASFV in hard ticks and sheep. **A.** Lane 1-9 used DNA from *D. silvarum* and Lane 10-18 used DNA from sheep blood. **B.** Pig represents the 235-bp DNA segment from the ASFV reference genome (GenBank: AY261365.1). Tick represents the 235-bp DNA segment from *D. silvarum* amplified by PCR and sequenced by Sanger technology. Sheep represent the 235-bp DNA segment from sheep blood amplified by PCR and sequenced by Sanger technology. *D. silvarum* and sheep blood were collected from two different places in Xinjiang province of China.

Further analysis of two 235-bp sequences from *D. silvarum* and sheep showed an identity of 100% between them (**Figure 1B**). Using the ASFV reference genome (GenBank: AY261365.1), three single nucleotide mutations were detected at the positions of 38, 76 and 108 (C38T, C76T and A108C) on these two 235-bp sequences. As the 235-bp ASFV segment encodes 78 amino acids, C38T results in a single amino acid mutation at the position 66 (G66D). C38T suggested the existence of a new ASFV strain, which was different from all reported ASFV strains in NCBI GenBank database (version 197). As the *D. silvarum* ticks and the sheep blood were collected from two different places, this new strain could infected sheep and be transmitted by *D. silvarum* ticks. Using blast tools, the results showed that the new 235-bp segment in the new ASFV strain did not contain high similar regions with sequences from other viruses, bacteria or animal genomes. As this segment was highly specific to represent ASFV, our study still proved the existence of ASFV in *D. nuttalli, D. silvarum* and sheep.

## Conclusion and discussion

In this study, we detected a new 235-bp ASFV segment in hard ticks from sheep and bovine. Although this segment was highly specific to represent ASFV based on knowledge from the current public databases, it still can not rule out the possibilities of this segment from unknown species. The further work is to obtain the complete genome sequence of ASFV from hard ticks or sheep.

## Materials and Methods

*A. testudinarium* ticks were captured from buffalo in Yunnan province of China, and *D. nuttalli* and *D. silvarum* ticks were captured from sheep and bovine in Xinjiang province of China. All ticks were used to pool four samples representing *A. testudinarium* adults, *D. nuttalli* adults, *D. nuttalli* larvae and *D. nuttalli* nymphs. Four samples were used to construct four sRNA-seq libraries to be sequenced using Illumina sequencing technologies with the length of 50 bp, respectively [11]. As the library of *D. nuttalli* adults was sequenced twice, five runs of sRNA-seq data were deposited at NCBI SRA database under the project accession number SRP084097.

The cleaning and quality control of sRNA-seq data were performed using the pipeline Fastq_clean [12] that was optimized to clean the raw reads from Illumina platforms. The virus detection was performed using the pipeline VirusDetect [6]. Statistical computation and plotting were performed using the software R v2.15.3 with the Bioconductor packages [13]. The ASFV reference genome (GenBank: AY261365.1) was used for all the data analysis in this study.

The RNA extraction of *D. nuttalli* and cDNA synthesis were performed using the protocol published in our previous study [14]. The DNA extraction of *D. nuttalli, D. silvarum*, sheep and bovine was performed using the protocol published in our previous study [15]. PCR amplification of DNA and cDNA using ASFV specific primers GCAGAACTTTGATGGAAACTTA and TCCTCATCAACACCGAGATTGGCAC to produce a 235-bp DNA segment (**Figure 1B**). PCR reaction was performed by incubation at 95 °C for 10 min, followed by 40 PCR cycles (15 s at 95 °C, 30 s at 62 °C, and 30 s at 72 °C for each cycle) and a final extension at 72 °C for 7 min.

## Authors’ contributions

ZC and SG conceived this project. SG and HY supervised this project. XY collected and identified the ticks. XX, XJ and HJ performed experiments. SG and XX analyzed the data. WD curated the sequences and prepared all the figures, tables and additional files. SG and ZC drafted the main manuscript. GL and JL revised the manuscript. All authors have read and approved the manuscript.

